# Concerns on research history overview, distribution patterns, and potentially neglected ecological consequences of the invasive apple snail (*Pomacea sp.*): A review

**DOI:** 10.1101/2023.12.23.573227

**Authors:** Du Luo, Yuefei Li, Jie Li

## Abstract

Invasive species pose a threat to global ecosystems and biodiversity, with the apple snail standing out as one kind of the worst invasive species due to its exclusive expansion. In order to draw a conceptual framework of its invasion pattern from an overview, we endeavor to conduct a bibliometric analysis and systematic review to explore the existing knowledge on research history and subjective categorization of *Pomacea canaliculata* and *P. maculata*, and to uncover emerging trends of distribution and possible neglected impact on food webs. The research history reveals that the first international research publication was documented in 1958. However, a substantial increase in international publications occurred only after 2000. The adoption and effective implementation of a universally agreed upon management strategy may greatly enhance the progress of research on apple snail invasion. Notably, co-word analysis revealed that the content of each thematic cluster has a shift from basic biology to the monitoring of occurrences and environmental adaptation. Currently, the global distribution of apple snails has expanded to 39 countries primarily as a result of anthropogenic translocation. All infested countries, with the exception of Israel and Singapore, are major paddy rice producers. The northernmost distribution limit is Tyumen in Russia, while the southernmost limit is the Buenos Aires Province in Argentina. Biological control has emerged as a prominent technology for the prevention of apple snail infestations, leading to the identification of 98 non-target enemy species as potential control agents. A taxonomic diversity analysis revealed that these species encompassed a wide range of phylogenetic classifications, including 6 Phyla, 12 Classes, 33 Orders, 55 Families, and 82 Genera. This observation suggests that there is a substantial pool of animals from diverse phylogenetic levels that can prey upon apple snails. Given the high population density and extensive distribution of apple snails, there is a pressing need for further research to explore their potential ecological ramifications. The current understanding of the upward ecological impacts of apple snails remains inadequate. These findings provide valuable insights for future research endeavors and contribute to the effective management of invasive apple snail populations.

## 1. Introduction

Invasive species pose a significant threat to ecosystem function, biodiversity, and human well-being, and are also associated with substantial economic costs (Henry et al., 2023). Economic data from the United States indicate that the majority of these costs are attributed to resource damages, with agriculture being the most affected sector (Fantle-Lepczyk et al., 2022). The impacts of invasions are multifaceted, including changes in native species richness and abundance, alterations in native animal behavior, and modifications to trophic networks. International policy agreements can play a crucial role in mitigating the impact of invasive alien species (Diagne et al., 2021). Effective management strategies rely on accurate species distribution information and control measures. About 100 freshwater snail species have invaded non-native fields. Among these, apple snails (*Pomacea sp.*) are considered one of the most widely spread global invaders (Preston et al., 2022).

Apple snails (Ampullariidae) are a group of freshwater snails comprising over 150 species belonging to nine genera (Hayes et al., 2009). The genus *Pomacea* is naturally found in the New World, with one species native to Florida, USA, and the remaining species native to South and Central America, including the Caribbean (de Brito and Joshi, 2016). Their remarkable environmental adaptability and unique reproductive behavior have contributed to their successful establishment and expansion in various habitats. Unlike many other freshwater snails that lay eggs underwater in a transparent gelatinous mass, all species of *Pomacea* lay their eggs above the water’s surface. This adaptation has facilitated their population growth. The species within the genus *Pomacea* have become significant agricultural and environmental pests in numerous countries worldwide. As early as 2000, the golden apple snail (*Pomacea canaliculata*) was recognized as one of the most invasive species by the IUCN (Lowe et al., 2000). From an early stage, the introduction of apple snail to East Asia and Southeast Asia had caused substantial detrimental effects on rice production (Halwart, 1994). Among the apple snail species in the *Pomacea* genus, *P. canaliculata* and *Pomacea maculata* are the two most invasive species originally from South America (Joshi et al., 2017).

Enduring successive dispersion over a period of four decades, *P. canaliculata* has emerged as the dominant aquatic gastropod in numerous infested areas. This species is also recognized as a major pest in rice agriculture (de Brito and Joshi, 2016). Phylogeographic analysis has revealed that *P. canaliculata* and *P. maculata* exhibit significant genetic diversity and pronounced genetic differentiation among different countries (Liu et al., 2019). Moreover, taxonomic shortcuts within the Ampullariidae family, particularly in the genus *Pomacea*, have frequently hindered species delineation and identification (Hayes, 2021). Furthermore, *P. canaliculata* serves as an intermediate host for *Angiostrongylus cantonensis*, leading to the emergence of human eosinophilic meningitis epidemics (Wang et al., 2022). The invasive apple snails (*P. canaliculata* and *P. maculata*) pose a global threat to ecosystems and biodiversity, while also causing substantial damage to agricultural production and increasing the risk of transmitting epidemic diseases.

In order to achieve sustainable management of apple snails, biocontrol has been regarded as a substitute for synthetic molluscicides (Azmi et al., 2022). As one of the most severe invasive animals expanding rapidly, a substantial amount of research knowledge and data have been accumulated worldwide. Consequently, there is a pressing need to explore prominent research trends within the extensive, fragmented, and sometimes controversial body of literature. Additionally, it is crucial to establish a comprehensive understanding of the current distribution of apple snails globally and to investigate their possible overlooked impact on predators within natural food webs.

## 2. Research history and thematic categories

We conducted a search in the Web of Science (WoS) database, accessed on July 21, 2023, using the query terms ‘*Pomacea*’ in the fields of themes and title. The search yielded a total of 1171 publications related to themes of *Pomacea* and 545 publications with *Pomacea* mentioned in the title. Notably, there was a higher number of publications focused on *P. canaliculata* compared to *P. maculata*. The earliest international publication on this topic dates back to 1958. However, before the year 2000, only 10 publications were found that specifically addressed *Pomacea*-related themes. In Asia, apple snails have been introduced several decades ago as aquaculture organisms due to their potential as high-quality aquatic food resources (Naylor, 1996).

China has long been the largest producer of aquaculture food globally and has also held the top position in rice (*Oryza sativa*) production. In light of this, we conducted a comparative analysis of apple snail research publications in Chinese, collected from the CNKI database, and international publications in English, gathered from the Web of Science database (Fig. 1). The introduction of apple snails to mainland China occurred in 1981, with the first apple snail-related publication in Chinese recorded in 1982. Subsequently, a substantial number of Chinese scientific research articles on apple snails were published starting from 1985. In fact, a total of 2085 publications related to apple snails in Chinese were categorized in the MedaLink database. The number of publications on *Pomacea sp.* experienced a significant increase in 2006 and reached its peak in 2008. Analysis of data from WoS and CNKI revealed that the number of publications remained equivalent in 2016, after which the international publication count surpassed that of Chinese publications.

**Fig. 1.**
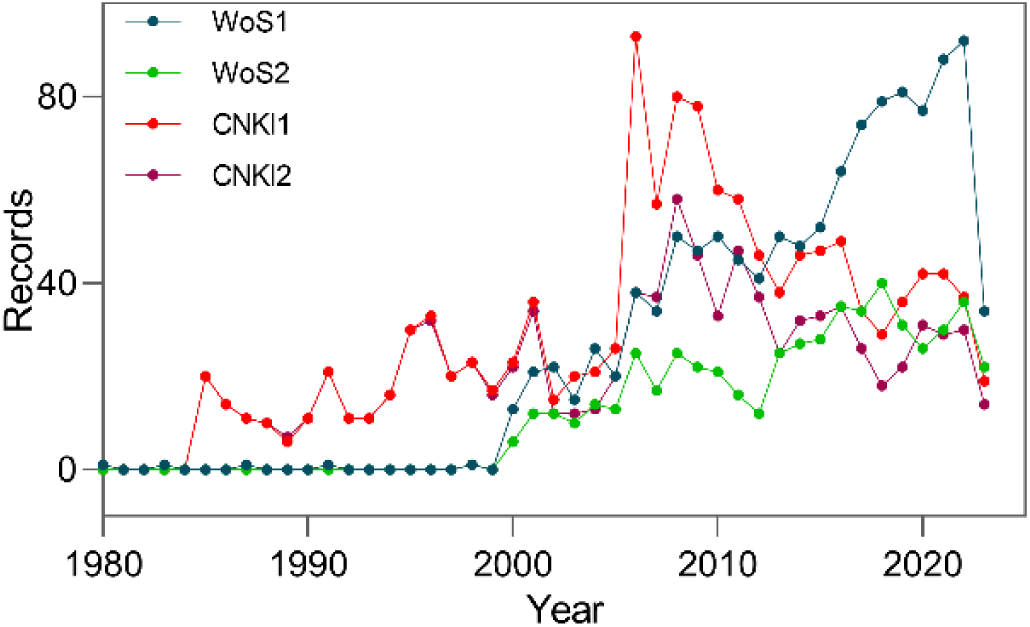
The figure depicts the count of scientific publications specifically focused on apple snails (*Pomacea sp.*), retrieved from the databases of Web of Science and CNKI. In the Web of Science database, the search was conducted using the terms “*Pomacea*” in the thematic fields (WoS1) as well as in the titles (WoS2) of the publications. Similarly, in the CNKI database, the search was performed using the Chinese translations of apple snails in the thematic fields (CNKI1) and both the titles and keywords (CNKI2). The timeframe for the search extended from 1980 to 2023.

The observed variations in publication numbers can be attributed to several key factors. Firstly, the designation of *P. canaliculata* as one of the most detrimental invasive species in 2000 (Lowe et al., 2000) likely influenced the subsequent research interest and publication output. Additionally, the emergence of the epidemic disease caused by *A. cantonensis* in China in 2006 (Lv et al., 2009) may have further contributed to the increased attention and subsequent publications on *Pomacea sp.* Furthermore, the continuous global expansion of apple snails has also played a role in driving research interest and publication trends up until the present time.

Tracing the disciplinary classifications, analysis of the searching results on themes of *Pomacea* from WoS reveal that previous studies predominantly focused on zoology, environmental ecology, and biology. Within the field of biology, investigations encompassed areas such as biochemistry, reproduction, development, and physiology. Additionally, research in agriculture explored aspects related to plant sciences, while studies in medicine primarily fell under the purview of life sciences. Among the various species, *P. canaliculata* and *P. maculata* emerged as the most extensively researched. Notably, population studies centered on biochemistry, molecular genetics, biogeography, and morphology emerged as prominent areas of interest, driven by the need to resolve taxonomical and identification questions surrounding apple snails. Furthermore, it is worth highlighting that approximately 70% of the papers with themes on *Pomacea* aligned with the field of environmental sciences ecology, signifying a significant international research interest in this domain.

In order to explore the intellectual structure of the specific contents, a bibliometric analysis was conducted using CiteSpace 6.1 R6, following the methodology outlined by Aria and Cuccurullo (2017) and Chen (2006). The aim was to synthesize existing knowledge through comprehensive science mapping analysis. To examine the actual content of publications on apple snail research, the technique of co-word analysis was primarily employed as described by Donthu et al. (2021). To obtain relevant literature, a search was performed in the Web of Science Core Collection (2000-present) using “*Pomacea*” as the keyword in both title and theme fields. Following removal of duplicate records, a total of 494 and 935 unique records were utilized for the bibliometric analysis, respectively. The results of the co-occurring keywords analysis provided insights into the latest trends and potential future directions of research, as suggested by Sabé et al. (2023) (Fig. 2).

**Fig. 2.**
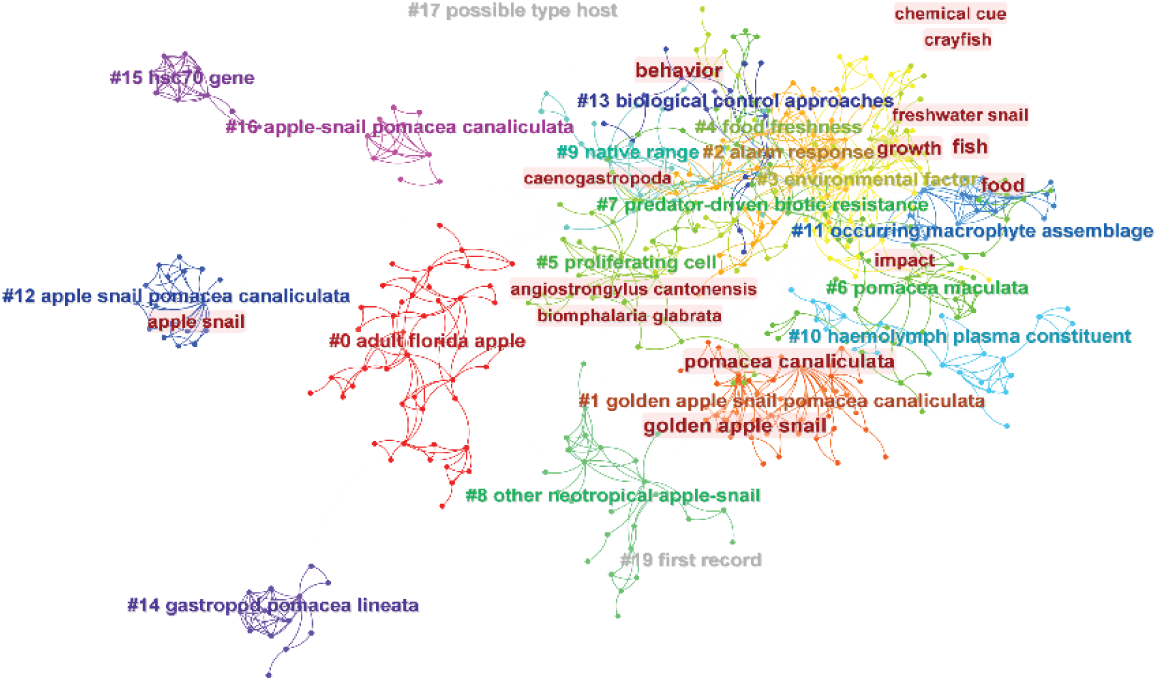
Illustrates the co-word (keyword co-occurrence) network using CiteSpace 6.1.R6. In this network visualization, each node represents a keyword, with the size of the node indicating the frequency of occurrence of the keyword. The links between nodes represent the co-occurrence relationships between keywords. Different thematic clusters are distinguished by different colors. Clusters labeled with “#” indicate evolved title terms. The data for this analysis were obtained from the Web of Science database (https://www.webofscience.com/) by querying the “title” field with the search terms “*Pomacea*”.

Burstness analysis of keywords revealed that the main research themes during the period from 2000 to 2010 were the biology of life history traits and herbivory. Around 2010, genetics and phylogenetic evolution emerged as a prominent subject category, which subsequently evolved into species identification and occurrence monitoring in recent years. In the field of environmental ecology, research focused on the tolerance of apple snails to environmental factors and their physiological-chemical activation, particularly in recent years. Since 2012, researchers have also begun to investigate the impact of apple snails on fisheries and biodiversity. Additionally, two other research areas concerning apple snails include their potential threat to human health and paddy rice production. However, the decline in research activity in these areas may be attributed to the lack of epidemic emergency reports and the increased use of chemical molluscicides in agriculture. Nevertheless, there are still some recent studies that explore the use of plant extracts for controlling apple snails.

In the Chinese language, we conducted a search in the CNKI database using three translations of “*Pomacea*” as “title or keywords”. A total of 937 records were retrieved for co-word analysis using CiteSpace 6.1.R6. The analysis revealed that during the late 1980s to early 1990s, the main research interests focused on the nutrition value and culturing technology of *Pomacea*. A significant increase in research attention towards the intermediate host burst was observed in 2007. Interestingly, researchers started showing more interest in preventing apple snail in 1993, approximately 12 years after its introduction, despite the existence of at least three scientific reports published as early as 1985, which highlighted the risks associated with culturing apple snails. In the field of environmental ecology, the primary focus of apple snail research shifted towards biological invasion since 2013. The latest research content mainly revolves around the following aspects: invasion monitoring combined with species identification and genetic analysis, adaptation and tolerance to important environmental factors, controlling apple snail using plant extracts and environmentally-friendly chemicals, and investigation on the intermediate host of *A. cantonensis*. To further advance apple snail research, there is a growing need to adopt interdisciplinary research methods, which can contribute to both quantitative and qualitative improvements in the field.

## 3. Global distribution pattern

### 3.1 Expansion history and current distribution with rice production

Although the release and escapes of *P. canaliculata* from aquariums and pet trade in the US may date back to the 1950s, its first discovery in North American waters occurred in 1978 in the county of Florida (Joshi and Sebastian, 2006). In Asia, apple snails (*P. canaliculata*) were intentionally introduced as high-quality food sources in 1979-1980. Subsequently, they rapidly invaded aquatic systems in Japan, the Philippines, China, Malaysia, Indonesia, and Thailand in the following years (Naylor, 1996). Though this non-native aquatic gastropod expanded exclusively in East Asia and Southeastern Asia as early as the 1980s, its first report in Europe occurred in 2009, specifically in the rice fields of the Ebro Delta in Spain (Castillo-Ruiz et al., 2018). In 2020, scientists from Kenya reported the establishment of an apple snail population (*P. canaliculata*) in the Mwea irrigation scheme, marking the first confirmed record of an established apple snail population in continental Africa (Buddie et al., 2021). Results from our literature survey revealed that apple snails (*P. canaliculata* and *P. maculata*) have undergone rapid expansion and are currently distributed in 39 countries worldwide (Fig. 3).

**Fig. 3.**
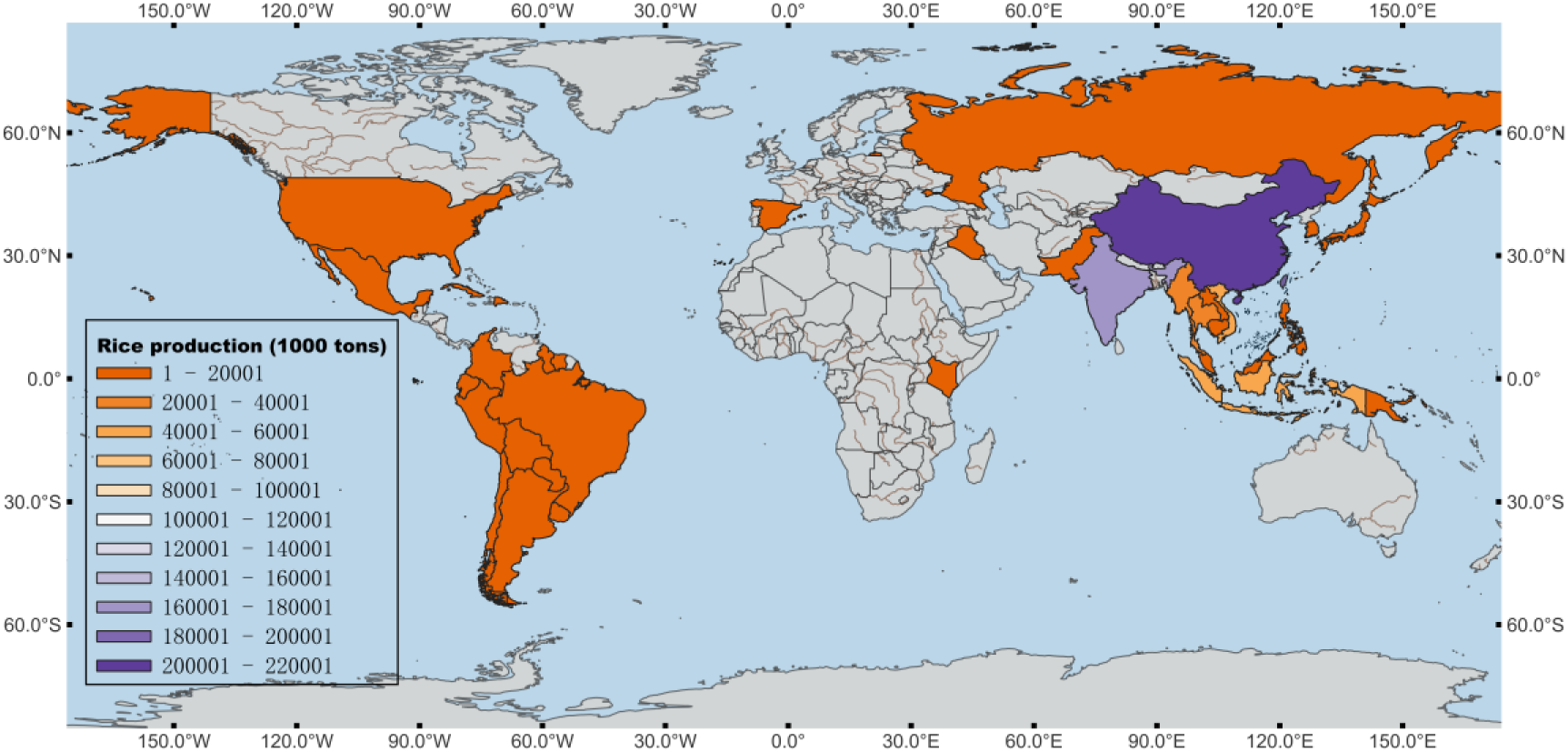
Global distribution of apple snail (P canaliculata and *P. maculata*). The map was generated using the free and open-source software QGIS. The legends on the map represent the gradient color for different countries, indicating paddy rice production (in thousand tonnes) based on data from the World Food and Agriculture Statistical Yearbook 2022.

Furthermore, according to existing records from GBIF (GBIF.org, 2023) and NAS-USGS (Benson, 2023), more scientific reports are still needed to assess the establishment of *P. canaliculata* and *P. maculata* in Guatemala, Panama, and Puerto Rico (Gilal and Muhamad, 2020). It is worth noting that, despite the absence of invasion reports, golden apple snails (*P. canaliculata*) were observed being sold in aquariums in Italy (Accorsi et al., 2014). Additionally, *P. canaliculata* has been sold as pets in South Africa (Shivambu et al., 2020) and Israel (Roll et al., 2009). In Switzerland, outbreaks of apple snails (*Pomacea sp.*) were under eradication in 2017, while in France, eradication efforts took place in 2018 (Schrader et al., 2020). Furthermore, *P. diffusa* was misidentified as P. bridgesii in Sri Lanka (Hayes et al., 2008), and *P. nobilis* was misidentified as *P. maculata* in Peru (Ramirez et al., 2020). Accurate identification of *Pomacea* species is crucial for their early detection and monitoring. It is important to note that climate change and introgressive hybridization may present new challenges in the monitoring and effective management of this highly invasive species (Yang et al., 2020).

In recent years, *P. canaliculata* has expanded its distribution range in native Argentina (Seuffert and Martín, 2021). The transportation by fishermen and aquarists is likely to be the main factor facilitating the establishment of populations (Seuffert and Martín, 2021). This extensive expansion may have overlooked impacts on diversity and ecosystem functions (Seuffert and Martín, 2021). Apple snails were predominantly found in sites near the shore with low current velocity and abundant organic matter (Seuffert and Martín, 2013). Only recently, the occurrence of apple snails was discovered in Kenya. However, subsequent investigations revealed that the paddy fields had been severely invaded. In a country-wide assessment, it was observed that all areas where apple snails had invaded were cultivating rice (*O. sativa*), with the exception of Israel and Singapore (Fig. 3). Asian countries account for 89.4% of rice production, and most of the infested countries in Asia are primary rice-production fields (FAO, 2022). Mainland India and Bangladesh, according to scientific reports, have been major paddy rice producers without apple snail coverage. In addition to earlier introductions for aquaculture purposes, aquarium hobby sales, as well as the transportation of aquatic vegetation, have been identified as the most common processes for introducing freshwater snails (Preston et al., 2022). The correlation between rice production and apple snail distribution requires further clarification. Monitoring and identification of *Pomacea* species in the aquarium trade may help forecast invasion risk.

### 3.2 Dispersion Limits at the Northernmost and Southernmost Extremes

Obviously, the long-distance expansion of the apple snail species was facilitated by anthropogenic translocation. In their native distribution, *P. canaliculata* had a southernmost range extension. However, *P. maculata* exhibited a larger range and extended northward to the south of the Amazon basin (Hayes et al., 2012; Seuffert and Martín, 2021). Notably, the expansion of *P. maculata* in southwest Louisiana occurred at a rate of 2-4 km per year (Lucero and Wilson, 2023). In China, a survey conducted during 2014-2015 revealed that natural populations of apple snails were distributed as far north as latitude 31.23°N in Jiangsu provinces (Yang et al., 2018). However, a more recent survey conducted in 2021 identified the invasion of both apple snail species as far north as Shandong province, with a location at latitude 35.61°N (Wang et al., 2022). In addition to these successful invasions, apple snails were found reproducing in two parks in Beijing in 2014 and 2020, respectively (Fan et al., 2021). Phylogenetic analysis suggests that the recent introduction of these species may have occurred through the transportation of aquatic plants from Zhejiang province. Even though the capability of overwintering may limit the establishment of apple snails, there are still seasonal invasion risks (Li et al., 2021). Outside the natural established areas in the south of China, apple snails have been observed in agricultural fields in eight provinces beyond the invasion areas between 2011 and 2020 (Zhuo et al., 2022). In this study, we investigated the global distribution limits of apple snails in both the southern and northern regions (Fig. 4). Our findings revealed that the southern limit of apple snails is located in the Buenos Aires province of Argentina (−38.42°N). Conversely, the northern limit is found in Tyumen of Western Siberia in Russia (57.15°N), where *P. canaliculata* individuals were sold as pets and subsequently introduced into aquatic ecosystems receiving thermal water from a nearby power plant (Vinarski et al., 2015). Additionally, in Twin Falls, Idaho, USA, *P. canaliculata* has stably established itself at a latitude of 42.66°N, facilitated by the discharge of warm water (Boler and Frest, 1991; Daniel, 2019; NAS sighting report). Moreover, in Japan and South Korea, apple snails have been established north of locations with a latitude higher than 35°N.

**Fig. 4.**
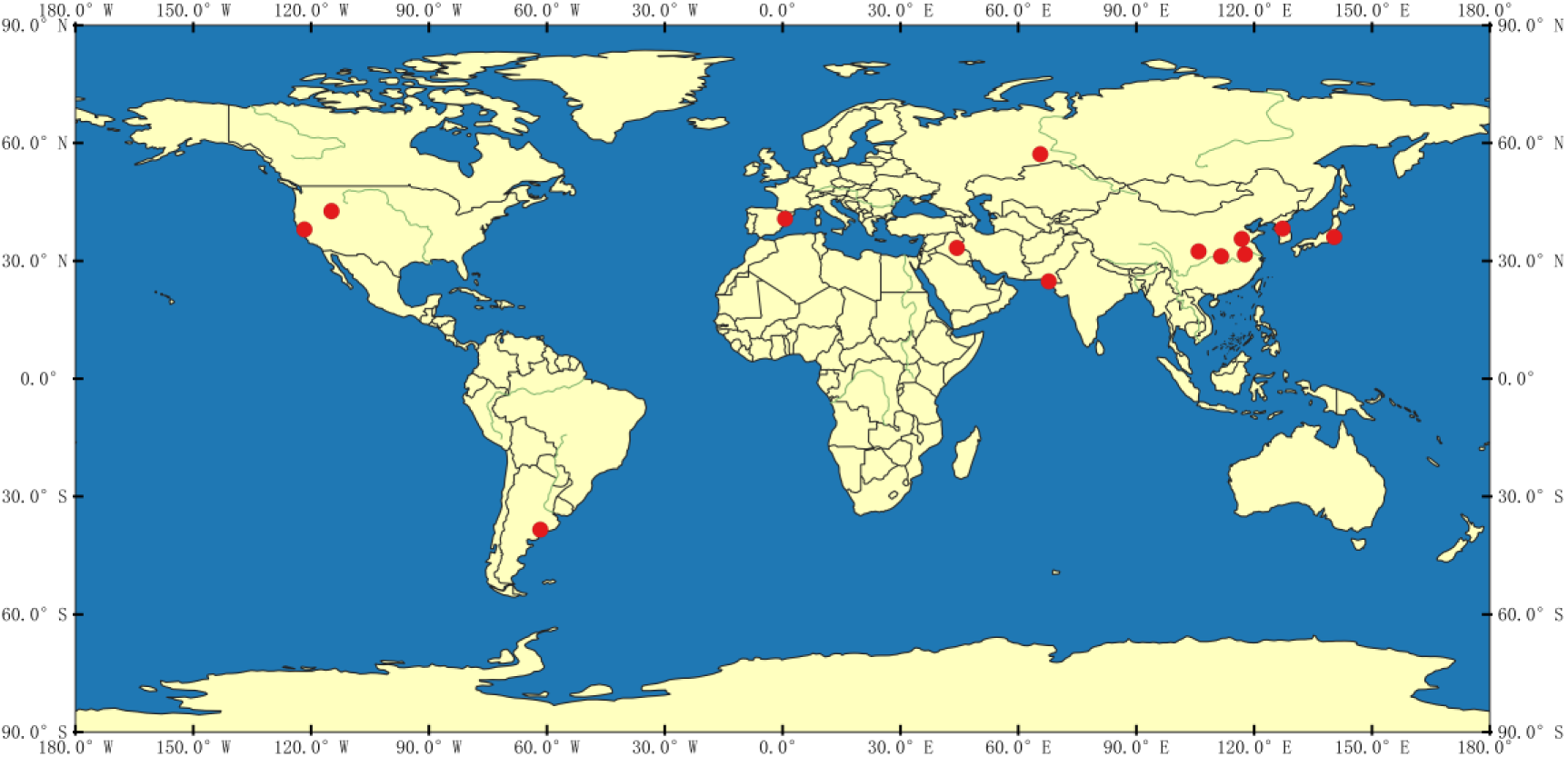
Global distribution of apple snails (*P. canaliculata* and *P. maculata*) along their northernmost limit and southernmost limit. The map was made with the free and open-source QGIS.

## 4. Neglected ecological impacts of non-target enemies in biocontrol

Controlling pests using natural enemies is a recognized ecosystem service in agriculture (Martin et al., 2013). Apple snails are widely acknowledged as significant pests in waterlogged cultivated areas. Among these, *P. canaliculata* has emerged as the dominant aquatic gastropod species causing damage to rice and toro crops across several Asian countries. In the realm of biological control methods against apple snails, a diverse range of organisms including animals, plants, fungi, and bacteria have been explored as potential natural enemies. Notably, microbial pathogens such as *Pseudomonas aeruginosa* and *Pseudomonas fluorescens* have demonstrated the ability to effectively eliminate *P. canaliculata* (Chobchuenchom and Bhumiratana, 2003). Additionally, entomopathogenic fungi *Paecilomyces lilacinus* have exhibited high efficacy in controlling apple snails by targeting newly hatched juveniles (Monchan et al., 2009). Another approach involves the use of weeds, specifically *Ambrosia artemisiifolia*, which can suppress the growth of *P. canaliculata* in rice fields without causing any adverse effects on rice plants (Panda et al., 2021). Furthermore, plant extracts have shown promise as effective biological measures against apple snails. In Brazil, extensive testing involving 1426 plant species was conducted to identify potential vegetal-derived molluscicides (Baptista et al., 1994). While various organisms have been utilized in the control of *Pomacea* species, natural enemies of animals have gained substantial attention and witnessed increased research efforts in recent years.

We conducted a comprehensive analysis of animals utilized as natural enemies for controlling apple snails, based on data derived from laboratory trials and potential field applications. The results revealed the presence of 98 species that exhibited potential as natural non-target animals for apple snail control. Taxonomic diversity classification demonstrated that these species were distributed among 6 Phyla, 12 Classes, 33 Orders, 55 Families, and 82 Genera (Fig. 5). This suggests a substantial availability of non-target enemies across various phylogenetic levels. The primary mechanism of biological control appeared to be predation. Among the predators identified, common carp (*Cyprinus carpio*) and African catfish (*Clarias gariepinus*) were observed to preferentially consume small snails in rice fields. However, their application is limited by the crop’s growth stage and the water level in the field (Su Sin, 2006). The predation potential of ducks varied depending on species, and their adaptation to rice field conditions was influenced by body size (Teo, 2001). Other potential predators such as leeches, crabs, common carp, turtles, ducks, and rats were capable of attacking adult snails. However, the majority of these predators were primarily found in rivers or ponds (Yusa et al., 2006). While apple snails are considered agricultural pests, their natural habitats are often located in the wild areas surrounding rice paddy fields. Wetland environments serve as a source of water and food for snail populations, and their distribution can expand through rice field irrigation and heavy rainfall events (de Brito and Joshi, 2016).

**Fig. 5.**
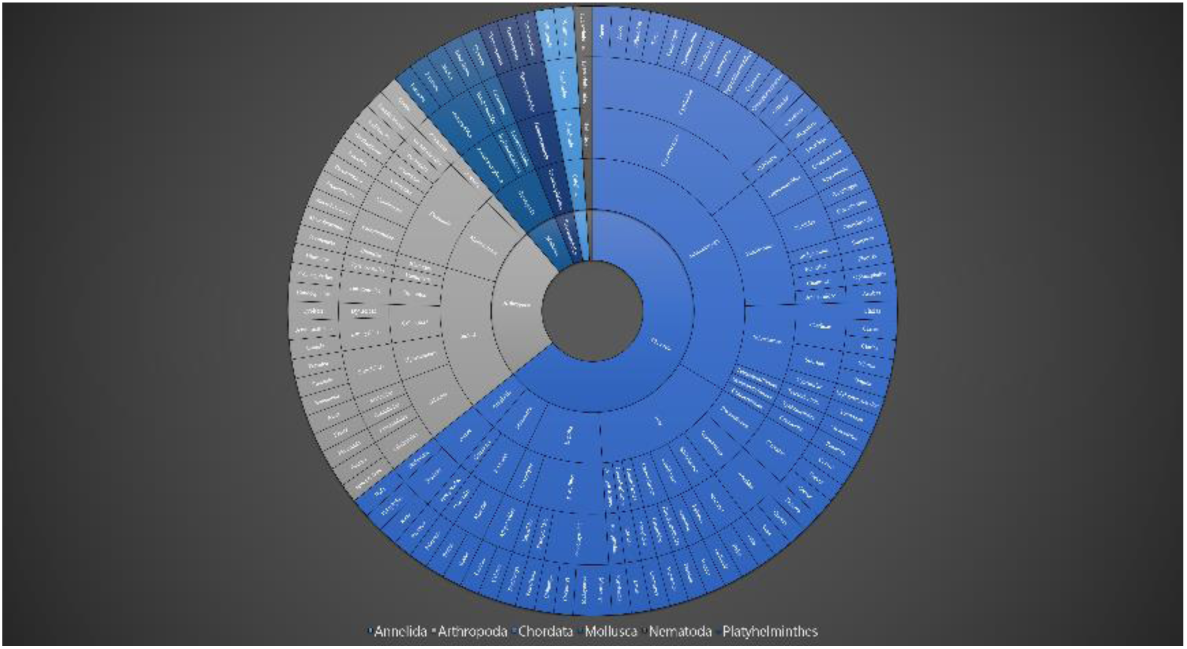
Taxonomic classification of non-target predators of apple snails (*P. canaliculata* and *P. maculata*) at different hierarchical levels. The layers in the doughnut chart, from the innermost to the outermost, represent Phylum, Class, Order, Family, and Genus.

Abundant historical research has focused on the impact of apple snails on ecosystems and the environment. Although apple snails are omnivores, plants constitute their main diet. Apple snails may selectively graze on wild macrophytes based on differences in nutritional value, chemical composition, and physical defenses, resulting in changes in floral diversity and functioning in wetlands (Qiu and Kwong, 2009). Additionally, their foraging behavior is size-dependent, and snails of all sizes have negative effects on the biomass of macrophytes and periphytic algae (Carlsson and Brönmark, 2006). Analysis of gut content from *P. canaliculata* in Singapore Quarry revealed that the snails’ diet consists of amorphous detritus, macrophytes, dinoflagellates, green algae, cyanobacteria, and invertebrate parts in varying proportions (Phoa, 2014). Stomach-content analysis also found that apple snails have a variable diet primarily consisting of detritus and macrophytes (Kwong et al., 2010). As an invasive herbivore, the impact of apple snails on the integrity and functioning of ecosystems has been a major research topic for many years. In fact, as early as 2004, a survey of natural wetlands demonstrated that high densities of snails can trigger a complete shift in ecosystem state and function (Carlsson et al., 2004). While there has been abundant research examining the direct downward effects of invasive apple snails, the indirect effects of their invasion remain insufficiently studied.

Based on the foraging ecology, the consumption of apple snails as prey would inevitably affect the growth, reproduction, and population dynamics of predators. Snails constitute a significant portion of the diet for certain species, including kites (*Rostrhamus sociabilis* and *Aramus guarauna*), lizards (*Dracaena guianensis*), and fish (Cowie, 2006). *P. canaliculata* has the ability to recognize predators, and the risk of predation has been shown to affect its growth and reproduction (Guo et al., 2017; Kahori and Yoichi, 2010). The anti-predatory behavior and alarm response further strengthen the predator-prey relationship in the biocontrol of apple snails (Ichinose, 2002; Xu et al., 2014). Native predators can rapidly develop a new preference for invasive aquatic mollusks as prey (Alexander et al., 2022). Even in newly invaded fields and wetlands in Europe, *P. maculata* has become a main prey item for native avian predators, such as *Plegadis falcinellus* (Bertolero and Navarro, 2018). The density of snails has been found to be positively correlated with key components of snail kite breeding biology (Cattau et al., 2014). Florida apple snails (*P. paludosa*) serve as the main food source for the Florida snail kite, *R. sociabilis*, and many other predators in wetlands. Therefore, the preference for specific snails as prey might impact the distribution pattern of their predators (Han et al., 2023). Furthermore, fish predators can indirectly impact the availability of prey for avian predators by altering the behavior and physiology of snails within the wetland food web (Siegfried et al., 2022). Even long-lived predators may respond rapidly to the invasion of apple snails (*P. maculata*) through morphological changes driven by phenotypic plasticity (Cattau et al., 2018). The estimated annual production of apple snails could be greater than any other freshwater snail populations, as indicated by published references (Kwong et al., 2010). Considering the high population density of apple snails in their microhabitat and their broad distribution worldwide, research on their potential ecological impacts remains insufficient.

## 5. Conclusion

The apple snail, known as a globally invasive species, poses risks to ecosystems, agricultural production, and human health. Our systematic review reveals that research on apple snails has been documented since the 1950s, with a significant increase in publications coinciding with the expansion of their invasion. Bibliometric analysis indicates a shift in research focus from basic biology to monitoring their occurrences and understanding their environmental adaptation. Through anthropogenic translocation, two invasive apple snail species, *P. canaliculata* and *P. maculata*, have established populations in 39 countries, most of which are rice (*Oryza sativa*) producing regions. It is worth noting that these snails have their northernmost distribution limits even reached areas beyond the subtropics. However, previous research on the ecological impact of these snails may be insufficient. Furthermore, evidence from studies involving non-target organisms used in biological control strongly suggests the existence of significant but overlooked ecological impacts. Therefore, future studies and management strategies should consider the high population density and extensive distribution of apple snails when exploring their potential ecological consequences.

## CRediT authorship contribution statement

Du Luo: Conceptualization, Data curation, Formal analysis, Methodology, Validation, Writing – original draft, Writing – review & editing. Yuefei Li: Conceptualization, Data curation. Jie Li: Data curation.

## Declaration of Competing Interest

The authors declare that they have no known competing financial interests or personal relationships that could have appeared to influence the work reported in this paper.

## Data availability

Data will be made available on request.

## Acknowledgements

This work was funded by the National Natural Science Foundation of China (Grant number 31600446).

## Notes

### Competing Interest Statement

The authors have declared no competing interest.

